# No evidence of early head circumference enlargements in children later diagnosed with autism in Israel

**DOI:** 10.1101/103788

**Authors:** Ilan Dinstein, Shlomi Haar, Shir Atsmon, Hen Schtaerman

## Abstract

**Background:** Large controversy exists regarding the potential existence and clinical significance of larger brain volumes in toddlers who later develop autism. Assessing this relationship is important for determining the clinical utility of early head circumference (HC) measures and for assessing the validity of the early overgrowth hypothesis of autism, which suggests that early accelerated brain development may be a hallmark of the disorder

**Method:** We performed a retrospective comparison of HC, height, and weight measurements between 66 toddlers who were later diagnosed with autism and 66 matched controls. These toddlers represent an unbiased regional sample from a single health service provider in the southern district of Israel. Using 4-12 measurements between birth and the age of two, we were able to characterize individual HC, height, and weight development with high precision and fit a negative exponential growth model to the data of each toddler with exceptional accuracy

**Results:** The analyses revealed that HC sizes and growth rates were not significantly larger in toddlers with autism even when stratifying the autism group based on verbal capabilities at the time of diagnosis. In addition, there were no significant correlations between ADOS scores at the time of diagnosis and HC at any time-point during the first two years of life

**Conclusions:** These negative results add to accumulating evidence, which suggest that brain volume is not necessarily larger in toddlers who develop autism. While early brain overgrowth may characterize specific individuals with autism, it is not likely to represent a common etiology of the entire autism population

## Background

Early brain overgrowth is one of the most prominent contemporary theories regarding autism development [1–3]. According to the theory autism may be caused by different genetic predispositions and/or environmental insults that accelerate cellular proliferation, migration, differentiation, and development so as to generate abnormally large brains during the first two years of life. Early accelerated growth is thought to be followed by arrested growth, which explains why adolescents and adults with autism do not have larger brain volumes [4]. Nevertheless, the availability of HC as a biomarker during early development could be extremely useful for identifying toddlers at risk of developing autism even before the onset of behavioral symptoms [2]. Since HC is an excellent correlate of brain volume during the first years of life [5], testing whether toddlers with autism indeed exhibit early brain overgrowth is a relatively straight forward venture

Previous studies have reported that toddlers who later develop autism are born with normal HC and then develop abnormally large head circumferences during the first two years of life [6–10]. These findings have been corroborated by postmortem [11] and MRI [12,13] studies that have reported significantly larger brains in toddlers who develop autism. Later studies, however, have questioned whether early overgrowth is specific to the brain or reflects general body overgrowth as apparent also in height and weight measurements [14]. Furthermore, the validity of early findings that were mostly based on comparisons with CDC population norms have been questioned, because these norms have under-estimated the true population HC [15]. With this in mind, several comparisons of early HC measurements between toddlers with autism and typically developing controls from the same community did not find any significant between-group differences [16–20]. These studies demonstrate the ongoing controversy regarding the existence and significance of early brain overgrowth in toddlers with autism.

When interpreting the studies above it is important to consider the sample characteristics in each case. For example, it has been suggested that larger brain volumes may be more strongly associated with specific autism etiologies involving regression [21], immunological insults [22], and specific genetic abnormalities (e.g., PTEN mutations [23]). Furthermore, HC is hereditary regardless of autism [24]. Differences in genetics and the environmental exposures of the examined sample as well as the inclusion/exclusion criteria of each study may, therefore, have an impact on potential differences across autism and control groups. For example, contrary to previous reports from the U. S. [25], assessment of large clinical databases in Norway and Israel reported modest [19] or no [26] differences in the rates of macrocephaly in children diagnosed with autism.

To further address these issues we performed a retrospective assessment of HC measurements that were recorded between birth and the age of two years old at Maccabi Healthcare infant wellness centers in the southern district of Israel. The collected data included 4-12 HC, weight, and height measurements from each of the 66 children who were later diagnosed with autism and the 66 community matched controls. This enabled us to examine HC development with high precision and compare longitudinal HC changes across groups while also fitting a growth model to the data of each toddler. In addition, we examined the potential relationship between HC and autism severity measures at the time of diagnosis as assessed by an autism diagnostic observation schedule (ADOS) exam [27]. Importantly, the examined data represents an unbiased community sample of the members of Maccabi Healthcare services who make up approximately one third of the population in southern Israel.

## Methods

### Participants

We collected retrospective HC, weight, and height data from electronic patient records of children who were diagnosed with autism at the Maccabi child development center in Beer Sheva (n=66, 60 boys). The autism group included children who were formally diagnosed with autism, Asperger's syndrome, or PDD-NOS between 2009 and 2015. All children were tested with the first [27] or second edition of the ADOS and the mean age of diagnosis was 36.94 months (13.05 SD). Each child with autism was matched, in terms of gender and date of birth (+/- 30 days), with a typically developing control child from the same southern Maccabi Healthcare district (Table 1). Children with known chromosomal disorders, hydrocephalus, additional developmental and neurological disorders, and those born before 36 weeks of gestation or below 2 kg were excluded from the study.

**Table 1:** Sample characteristics including the number of toddlers (relative number of females), the term in weeks, ADOS and ADI scores, age at diagnosis, and number of HC samples available from the individual patient records. Range of the sample is in parentheses.

|  | Autism | Control |
| --- | --- | --- |
| Number of toddlers (females) | 66 (6) | 66 (6) |
| Term in weeks (range) | 39.1 (36-42) | 39.5 (36.2-41.5) |
| ADOS total (range) | 16.1 (7-26) |  |
| ADOS severity score (range) | 7 (4-10) |  |
| ADI-R (range) | 18 (7-26) |  |
| Age at diagnosis (range) | 3 (1-5.3) years |  |
| Number of HC samples | 8.3 (4-12) | 8.4 (4-12) |

### Head Circumference, Weight and Height Data

We extracted all of the available HC, weight, and height measurements from the electronic patient records of each child. The majority of measurements were performed by nurses at Maccabi Healthcare Baby Clinics. Parents usually arrive at the baby clinic with documentation of an initial HC and weight measurement from the time of birth, which is copied into the child’s electronic records. The mean number of measurements per child was 8.3 (SD: 2.1) in the ASD group and 8.42 (SD: 1.74) in the control group. All participating children had at least four HC measurements. In Israel, all children are invited to visit a baby clinic at the ages of 1, 2, 4, 6, 9, 12, 15, 18, and 24 months to complete standard check-ups and immunizations as part of the free social health care services. We, therefore, focused our analyses on the first two years of life where retrospective HC, weight, and height measurements are abundant.

### Data Analyses

Analyses were performed with custom written code in Matlab (*Mathworks Inc.,USA*). Since each child had measurements at different time-points within the first 24 months of life, we first linearly interpolated the measurements into vectors with a time resolution of weeks. This enabled us to estimate the week-by-week HC, weight, and height values of each child from the age of their first measurement to the age of their last measurement (Figure 1). We then compared HC, weight, and height of children who later developed autism and controls while binning the data into three month intervals from birth to the age of 24 months. We used two-tailed two-sample t-tests to assess the significance of differences across groups. We also computed the Pearson’s correlation coefficient between ADOS severity scores (see below) and HC measurements for each of the age intervals (bins) described above. The significance of the correlation coefficients was not corrected for multiple comparisons in order to increase sensitivity.

ADOS Calibrated Severity Scores (also known as ‘Comparison Score’) were calculated using the ADOS-2 algorithm [28] in children who were diagnosed by the ADOS-2 (n=44) or an alternative algorithm [29] available for children who were diagnosed with the previous version of ADOS (n=22).

We used a negative exponential growth model that was used in previous HC studies of autism [16] to estimate the rate of HC growth in each toddler during the first two years of life:

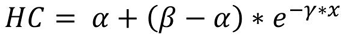

The model describes a nonlinear function where *α* represents the asymptote (a maximum size for growth within the time-frame considered), *β* represents the intercept at age x = 0 (i.e., birth), and γ represents the anti-log of the rate of change (how rapidly or slowly growth occurs from birth to the asymptote). The 3 growth parameters were estimated for each of the toddlers who had at least 5 HC measurements (i.e., 60 toddlers with autism and their matched controls). This was done to avoid potential over-fitting of the model in subjects with few data points. Performing the same analysis, however, with all toddlers yielded equivalent results. We used the same model to examine weight and height growth as well. We tested for group differences by performing two-tailed t-tests with each of the model parameters. The significance of differences across groups was not corrected for multiple comparisons in order to increase sensitivity.

## Results

### Individual development curves

We examined the developmental changes in HC, weight, and height, of individual toddlers from each of the groups during the first two years of life. All three measures exhibited the typical logarithmic shape and the distribution of individual values was qualitatively similar across the two groups (Figure 1). Note the high-resolution of these linearly interpolated growth curves, which is a feature of the large number of samples obtained from each toddler (see Methods).

**Figure 1:**
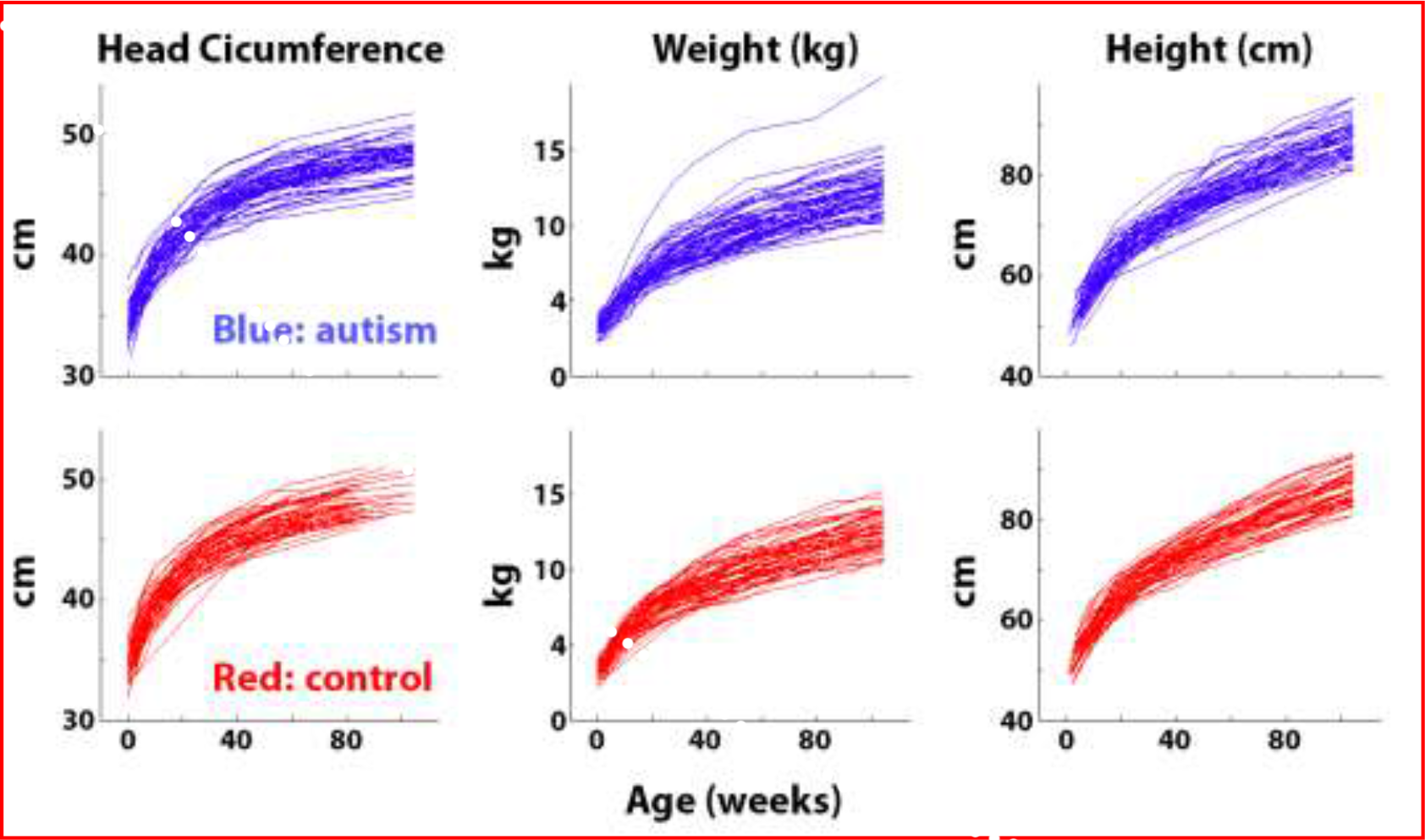
Head circumference, weight, and height measurements from the 66toddlers who later developed autism (blue) and the 66 matched controls (red). Each line represents the development of a single toddler as estimated with linear interpolation between measurement points (see methods).

### Comparisons across autism and control groups

Mean growth curves of the two groups were overlapping for all three measures (Figure 2, top row). We tested for differences across groups at multiple ages between birth and 24 months in three month intervals. The control group exhibited significantly larger head circumference and weight at birth and 3 months of age (p<0.05, t>2.1, uncorrected to increase sensitivity, Figure 2, bottom row). All other between-group comparisons did not reveal any statistically significant differences across groups.

**Figure 2:**
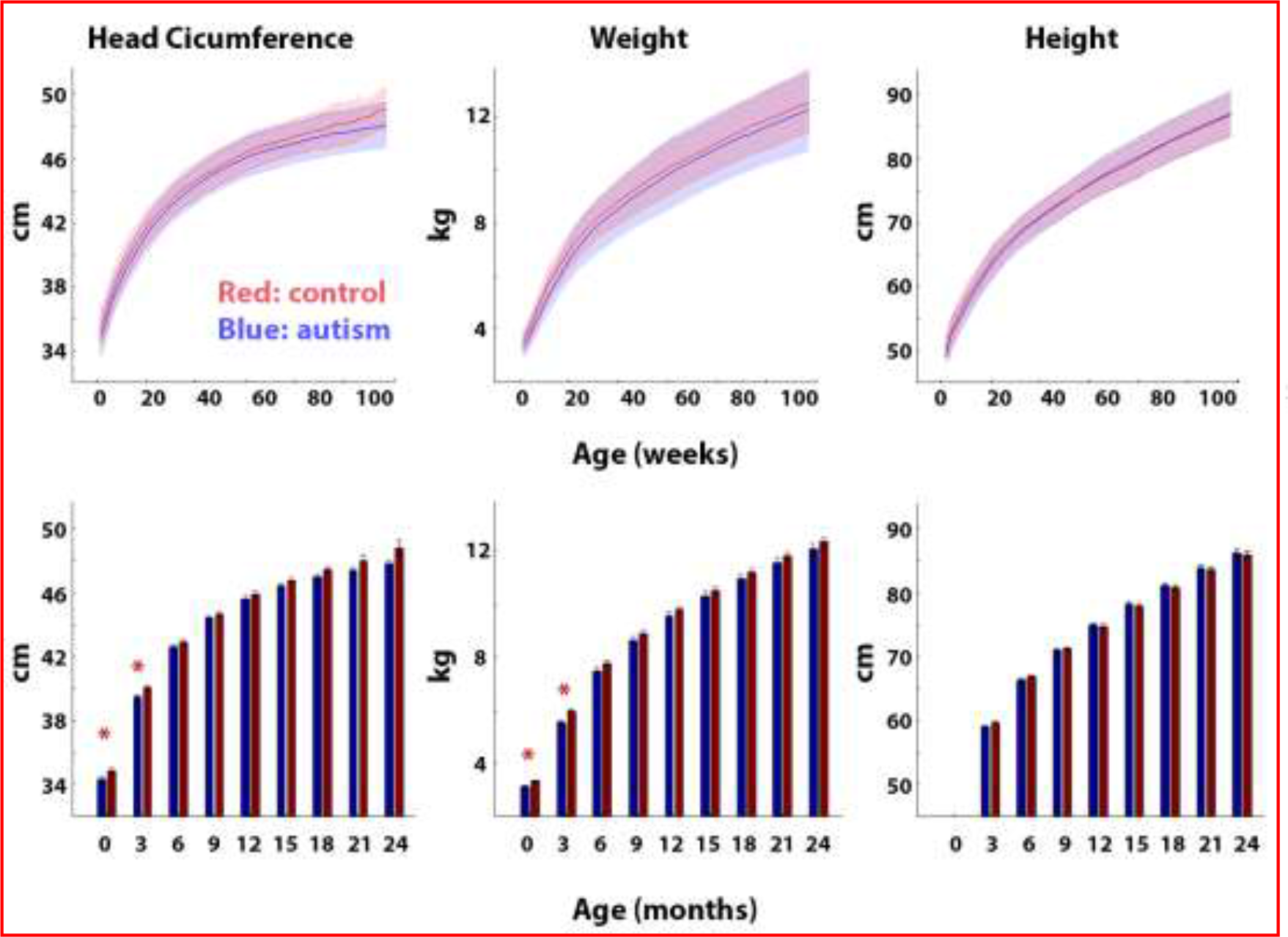
Comparison of HC, Weight, and Height across autism (blue) and control (red) groups. **Top row**: Mean of each group as a function of age in weeks. Shaded area: standard deviation across toddlers. **Bottom row**: Mean of each group in three month periods. Error bars: Standard error of the mean across subjects. Red asterisks: significantly larger values in the control group (p<0.05, uncorrected to increase sensitivity).

### Comparison of growth model parameters

We fit a nonlinear growth model (see Methods), estimating 3 growth parameters: intercept, rate of change, and asymptote for each toddler who had at least five HC measurements (i.e., 60 toddlers with autism and their matched controls, Figure 3). In agreement with the results described above, control toddlers exhibited significantly larger intercept (*β*) values when modelling HC or weight measures (p<0.02, t>2.3, Figure 3 B&C). This indicates that HC and weight measures were significantly larger in control toddlers as also described above (Figure 2). All other parameters did not differ across groups for any of the measures.

Note that the model fit the data of individual toddlers extremely well for all three measures as demonstrated by the scatter plots of individual R^2^ values. Performing the same analysis with all of the toddlers or only with males produced equivalent results: control toddlers exhibited significantly larger intercept (*β*) values when modelling HC or weight measures (p<0.03, t>2.2 in all cases).

**Figure 3:**
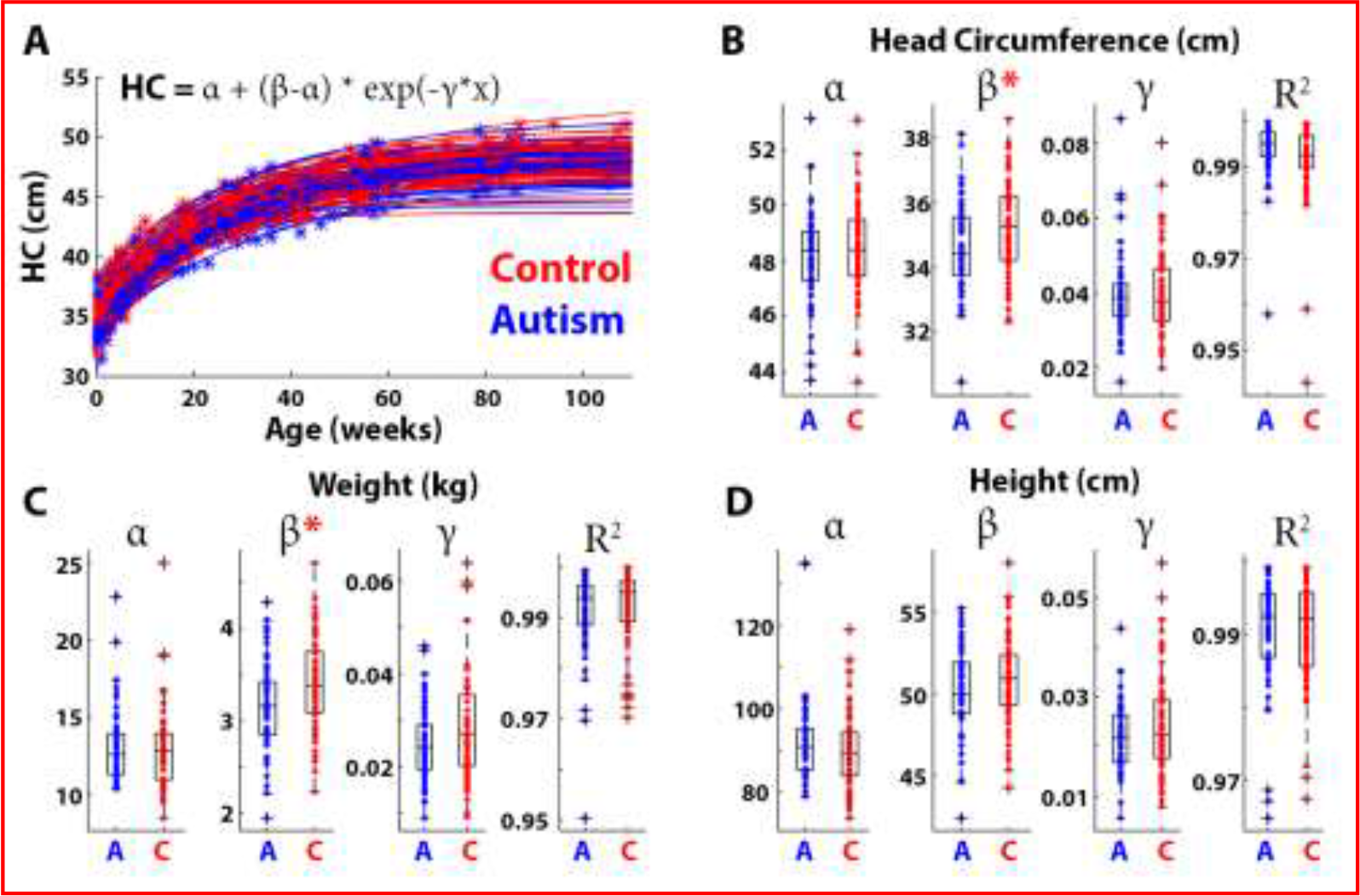
Comparison of head growth modeling parameters across autism (blue) and control (red) groups. Demonstration of HC growth model fits in individual toddlers: asterisks represent measurements and the lines represent the model fits (**A**). Boxplots demonstrating the HC (**B**), Weight (**C**), and Height (**D**) growth parameters: asymptote (*α*), intercept (*β*), and rate of change 
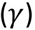
 in each group. Red asterisks: significantly larger parameter values in the control group. R^2^: Box plots demonstrating the model fits of individual toddlers in each group for each of the measures.

### Head circumference and autism severity

We examined the potential relationships between HC as measured at different ages and autism severity as quantified by the ADOS total score or calibrated severity score (see Methods) at the time of diagnosis. None of the correlations were statistically significant, but note that the correlation between HC and ADOS scores becomes positive and larger when using HC measures from older ages (Figure 4).

**Figure 4:**
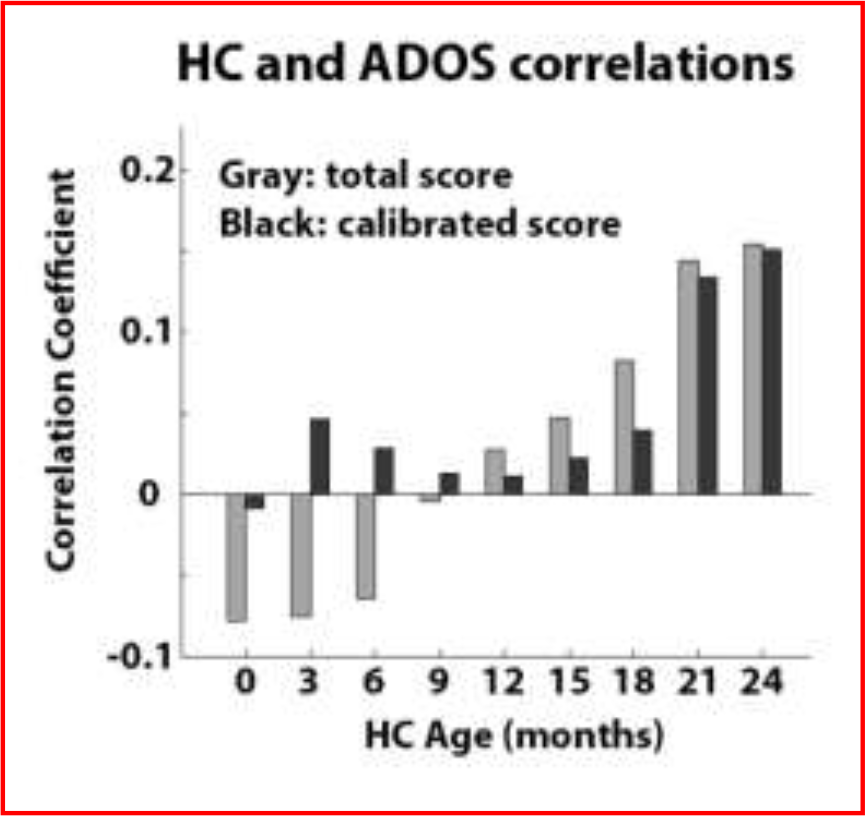
Relationship between HC at specific ages and ADOS score at time of diagnosis. Pearson’s correlation coefficients were computed for HC and total ADOS (gray) scores or calibrated ADOS scores (black).

### Verbal and non-verbal toddlers

We performed the same HC comparisons across groups while separating non-verbal toddlers who were diagnosed with the toddler module (n=14) or module 1 (n=36) from the verbal toddlers who were diagnosed with module 2 (n=15) of the ADOS test. Significantly larger HC was apparent in the control group at 3 months of age in comparison to the autism toddlers diagnosed with module 1 and at 24 months of age in comparison to the autism toddlers diagnosed with module 2 (p<0.05, t > 2.4, two-tailed t-test, uncorrected to increase sensitivity). All other differences between autism and control groups were not significant (Figure 5).

**Figure 5:**
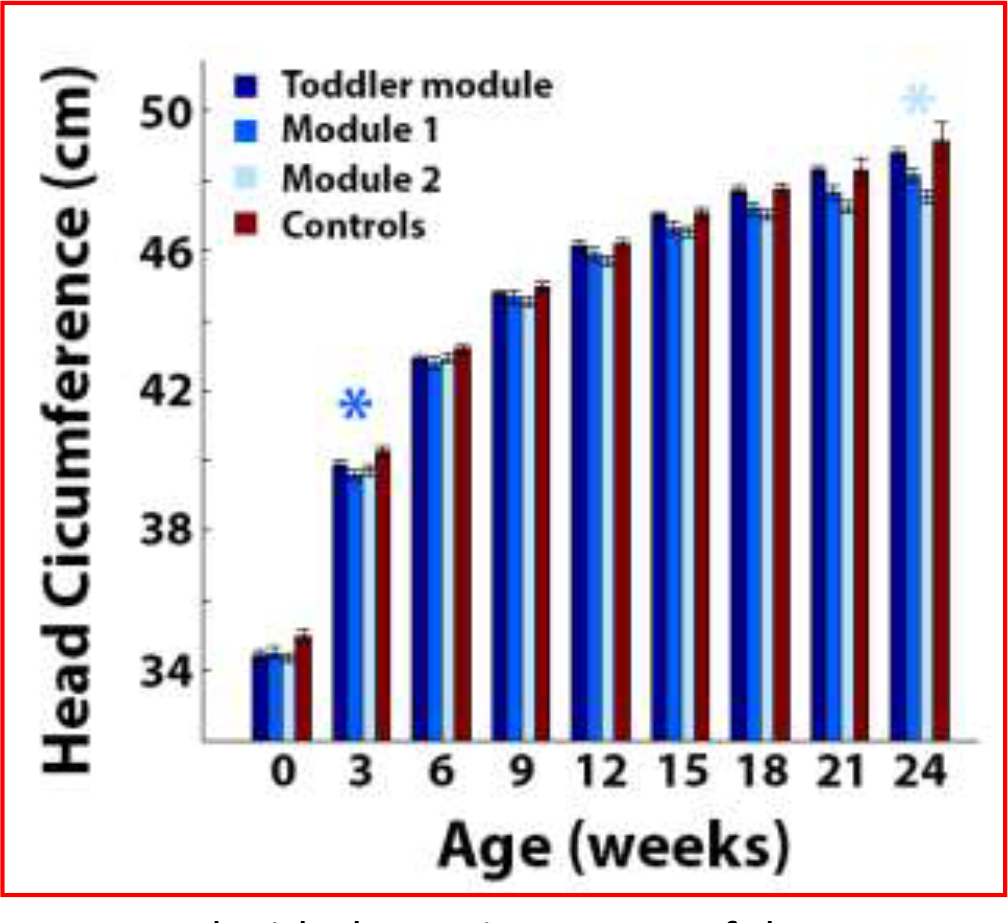
Comparison of HC at specific ages when separating the autism group according to ADOS modules. Mean HC in toddlers with autism diagnosed with the toddler module (dark blue), module 1 (medium blue), or module 2 (light blue) and control toddlers (red). Error bars: standard error of the mean across toddlers in each group. Asterisks: significantly larger HC in the control group as compared with the autism group of the corresponding color (p<0.05, uncorrected to increase sensitivity).

## Discussion

Our results demonstrate that toddlers diagnosed with autism in southern Israel do not exhibit early HC overgrowth during the first two years of life. Comparisons with control toddlers who were born at equivalent times and geographical locations revealed that HC, weight, and height measurements were mostly indistinguishable from those of toddlers with autism during the first 24 months of development and, if anything, were larger in the control group at birth and 3 months of age (Figures 1-2). Furthermore, growth parameters estimated using a non-linear negative exponential model were not significantly different across groups except for the intercept parameter, which indicated significantly larger HC and weight measures in the control group at birth (Figure 3). Equivalent findings were evident when examining only male toddlers and when splitting toddlers with autism into verbal and non-verbal groups according to the ADOS modules used for diagnosis (Figure 5). Finally, HC during the first 24 months of development did not predict later autism severity at the time of diagnosis (Figure 4).

An important strength of the current study lies in the relatively large number of measurements that were extracted from each of our 132 participant (4-12 HC, weight, and height measurements). This enabled us to estimate HC, weight, and height growth rates with high accuracy as reflected in the goodness-of-fits of individual toddlers (Figure 3). Furthermore, each of the 66 toddlers with autism were closely matched to a control toddler who was born within 30 days in the same geographical location. This ensured that the examined HC, weight, and height measurements were collected by similarly trained clinical staff within the same HMO district. Hence, the growth rates presented in the current study are likely to represent the true distribution of rates in the autism and control populations of southern Israel.

### Early brain overgrowth in autism

Previous studies have reported that HC is enlarged in autism during the first two years of life [6–10,12,14]. While most of these studies were biased by the use of misleading population norms published by the CDC [15], some studies have reported significant differences even in comparisons to community controls [12,14]. In addition, several MRI [12,13,30] and post-mortem [11] studies have reported that toddlers with autism exhibit larger and heavier brains than those of matched controls. This evidence forms the basis for the prominent early overgrowth theory of autism, which suggests that autism may be caused by abnormal cellular proliferation, migration, and differentiation that generate accelerated brain growth during the first two years of life [1–3].

With that said a growing body of evidence has reported that HC measures are not significantly larger in toddlers with autism when these are matched with control toddlers from the same community [16–20]. Our results are in line with these later studies and present further evidence for a lack of HC difference across groups in terms of both absolute HC measures and relative HC growth rates during the first two years of life.

When considering the conflicting results of the studies above it is important to remember that many of the earlier studies were based on small samples of 20-30 individuals in each group. Given the large heterogeneity in HC across individuals, significant differences across groups can be generated by a small number of participants with extreme measures. This is particularly relevant given the large variability in macrocephaly rates across different autism studies [25] and the existence of syndromal subtypes of autism that are specifically associated with macrocephaly (e.g., PTEN mutations [23]). While older, smaller studies in France and the U. S. have estimated macrocephaly rates in autism at approximately 15% [31,32], recent assessments of large clinical databases in Norway and Israel have reported more modest rates of 8.7% [19] and 4.4% [26] respectively. In addition, studies have reported that larger brain volumes are associated with specific autism etiologies involving regression [21] or immunological insults [22], while others have reported that HC is hereditary regardless of autism [24]. Differences in genetics and environmental exposures of specific samples as well as differences in inclusion and exclusion criteria across studies may, therefore, influence HC estimates.

## Conclusions

Given the large heterogeneity of mechanisms that have been implicated in the development of autism [33], it is not surprising that there are mixed reports regarding the existence of HC differences across autism and control groups in different studies. Indeed, similarly conflicting reports exist with respect to many physiological and behavioral measures studied in individuals with autism [34]. The great challenge facing contemporary autism research is to define specific sub-groups of toddlers with autism who exhibit specific etiologies. While early brain overgrowth may embody one such etiology, it is likely to be relevant to particular syndromic forms of autism and potentially to a small group of individuals who exhibit macrocephaly early in life. Future research regarding early overgrowth would, therefore, benefit from targeted studies with these particular individuals rather than attempting to associate early overgrowth with the entire autism population.

## Acknowledgements

This research was funded by an Israeli Science Foundation grant 961/14 (ID) and German Israeli Foundation grant I-2351-201.2/2014 (ID). We would like to thank the staff at the Maccabi Healthcare child development center in Beer Sheva for their help with data collection.

